# Molecular assembly of the KCNQ1–KCNE1–BACE1 complex

**DOI:** 10.64898/2026.03.03.709230

**Authors:** Alina Martin, Valeska Bienert, Stephanie Haefner, Florian Stockinger, Alexander Möhwald, Melanie Freimuth, Sandra Karch, Johannes Broichhagen, Guillaume Sandoz, Christian Alzheimer, Tobias Huth

## Abstract

We previously showed that the β-secretase BACE1 directly modulates KCNQ1 channels through a non-proteolytic mechanism. Here, we dissect the molecular interplay among KCNQ1, its canonical auxiliary subunit KCNE1, and BACE1 using bimolecular fluorescence complementation (BiFC), electrophysiology, single-molecule pull-down (SiMPull), and a Förster resonance energy transfer (FRET)-based interaction assay. BiFC validated KCNQ1 homotetramerization and confirmed specific interactions of KCNQ1 with KCNE1 and with BACE1 at the plasma membrane. To map interaction determinants, we generated six KCNE1/BACE1 chimeras. BiFC and electrophysiological recordings revealed domain-specific contributions: BACE1’s large extracellular domain primarily mediates its modulatory effects on KCNQ1 gating, while—consistent with previous reports—the KCNE1 transmembrane segment is necessary and sufficient to confer an *I_Ks_*-like phenotype, and the KCNE1 intracellular domain tunes the voltage dependence of activation. Notably, a chimera combining the BACE1 extracellular region with KCNE1 transmembrane and intracellular regions produced an *I_Ks_*-like current with additional BACE1-like slowing of activation, indicating functional additivity. Stoichiometry measurements by SiMPull and bleaching-step analysis demonstrated that KCNQ1 channel complexes predominantly recruit two BACE1 molecules. Although BACE1 oligomerizes at the plasma membrane, BiFC and FRET showed that KCNQ1 co-expression reduces BACE1 homomeric assembly, and this effect is unchanged by KCNE1 co-expression. Together, our data support a model in which BACE1 binds directly to KCNQ1, occupies a site distinct from KCNE1, and modulates KCNQ1 gating via its extracellular domain while remaining compatible with KCNE1 co-assembly.

## Introduction

Voltage-gated potassium channels of the KCNQ family are essential determinants of cellular excitability and potassium homeostasis. The KCNQ1 channel is expressed in a wide range of tissues such as the heart, inner ear, pancreas, kidney, and brain (Jespersen et al., 2005; Robbins, 2001). The KCNQ1 gene encodes the pore-forming α-subunit of the K_V_7.1 channel, which assembles as a homotetramer (Sun & MacKinnon, 2017, 2020). Its biophysical properties are profoundly shaped by accessory subunits, especially KCNE1-KCNE5 subunits, and other factors including phosphatidylinositol 4,5-bisphosphate (PIP_2_), calmodulin (CaM), protein kinase A (PKA) modulation, and adenosine triphosphate (ATP) (Wang et al., 2020; X. Wu & Larsson, 2020). In the heart, co-assembly with the single-pass transmembrane β-subunit KCNE1 converts the relatively fast-activating KCNQ1 current into the slowly activating delayed rectifier current *I_Ks_*, which is critical for timely action potential repolarization (Wang et al., 2020; J. Wu et al., 2016; X. Wu & Larsson, 2020). Beyond the heart, KCNQ1/KCNE1 complexes are expressed in the stria vascularis of the inner ear, where they contribute to generating the endocochlear potential (Nin et al., 2016), and in various tissues involved in transport processes and homeostasis (Dixit et al., 2020).

Mutations in either KCNQ1 or KCNE1 can impair *I_Ks_*, causing congenital long QT syndrome and, in some cases, deafness (Kekenes-Huskey et al., 2022; Vyas et al., 2016). Because of the physiological and clinical importance of this channel complex, a long-standing goal has been to elucidate how the pore-forming α-subunit interacts with its β-subunits, and to define the stoichiometry and arrangement of accessory subunits within the channel. While recent work supports a variable incorporation of up to four KCNE1 subunits per KCNQ1 tetramer (Wang et al., 2019; X. Wu & Larsson, 2020), the precise assembly rules and potential competition with other auxiliary proteins remain unresolved.

We recently identified the β-secretase BACE1—best known for its role in amyloid precursor protein cleavage in Alzheimer’s disease—as an additional KCNQ and KCNE channel-interacting protein (Agsten et al., 2015; Heininger et al., 2025; Hessler et al., 2015; Sachse et al., 2013). Notably, BACE1 modulates K_V_7.1 gating independently of its proteolytic activity and alters *I_Ks_* when co-expressed with KCNE1 (Agsten et al., 2015). This raises the intriguing possibility that BACE1 may compete with, complement, or otherwise influence KCNE1’s effects on KCNQ1 (Lehnert et al., 2016).

Here, we investigate the molecular interplay between BACE1 and the KCNQ1/KCNE1 channel complex. Using bimolecular fluorescence complementation (BiFC), Förster resonance energy transfer (FRET), single-molecule pull-down (SiMPull) assays, and a panel of KCNE1/BACE1 chimeric constructs, we map the domains responsible for binding and modulation. Whole-cell patch-clamp recordings reveal distinct and, in some cases, additive effects of BACE1 and KCNE1 on KCNQ1 gating. Stoichiometric analysis by SiMPull further shows that up to two BACE1 subunits can associate with a single KCNQ1 tetramer, and that KCNQ1 co-expression disrupts BACE1 multimerization. Together, these findings uncover a previously unrecognized layer of KCNQ1 regulation and suggest that BACE1’s non-proteolytic functions extend to shaping the behavior of critical cardiac and epithelial potassium channels.

## Methods

### Plasmids

The following constructs were used in this study: hKCNQ1 (NM_000218.2) in pcDNA3.1, hKCNE1 (NM_001127670.1) in pcDNA3.1, hBACE1 (NM_012104.4) in pcDNA3.1, hENaC1α (NM_001038.5) in pcDNA3.1, rKir2.1 in pcDNA3, and hAPP in pCEP4.

For BiFC assays, C-terminal fusions of V_N_ (Venus residues 1–210) or V_C_ (Venus residues 211–238) were generated by appending each Venus fragment to the gene of interest. V_N_ constructs contained a 10-glycine linker followed by a His tag, whereas V_C_ constructs contained a 10-glycine linker followed by a Flag tag. Chimeric KCNE1/BACE1 constructs were assembled in pcDNA3.1 as outlined in Table 1. To generate N-terminally HA-tagged constructs, hKCNQ1 and hBACE1 were subcloned into a pCMV-HA expression vector, resulting in fusion of an HA epitope to the N-terminus of each protein. For hBACE1-BBS, the bungarotoxin binding site (BBS; TGGAGATACTACGAGAGCTCCCTGGAGCCCTACCCT) was inserted immediately downstream of the hBACE1 pro-peptide (after residue 45).

**Table 1.**
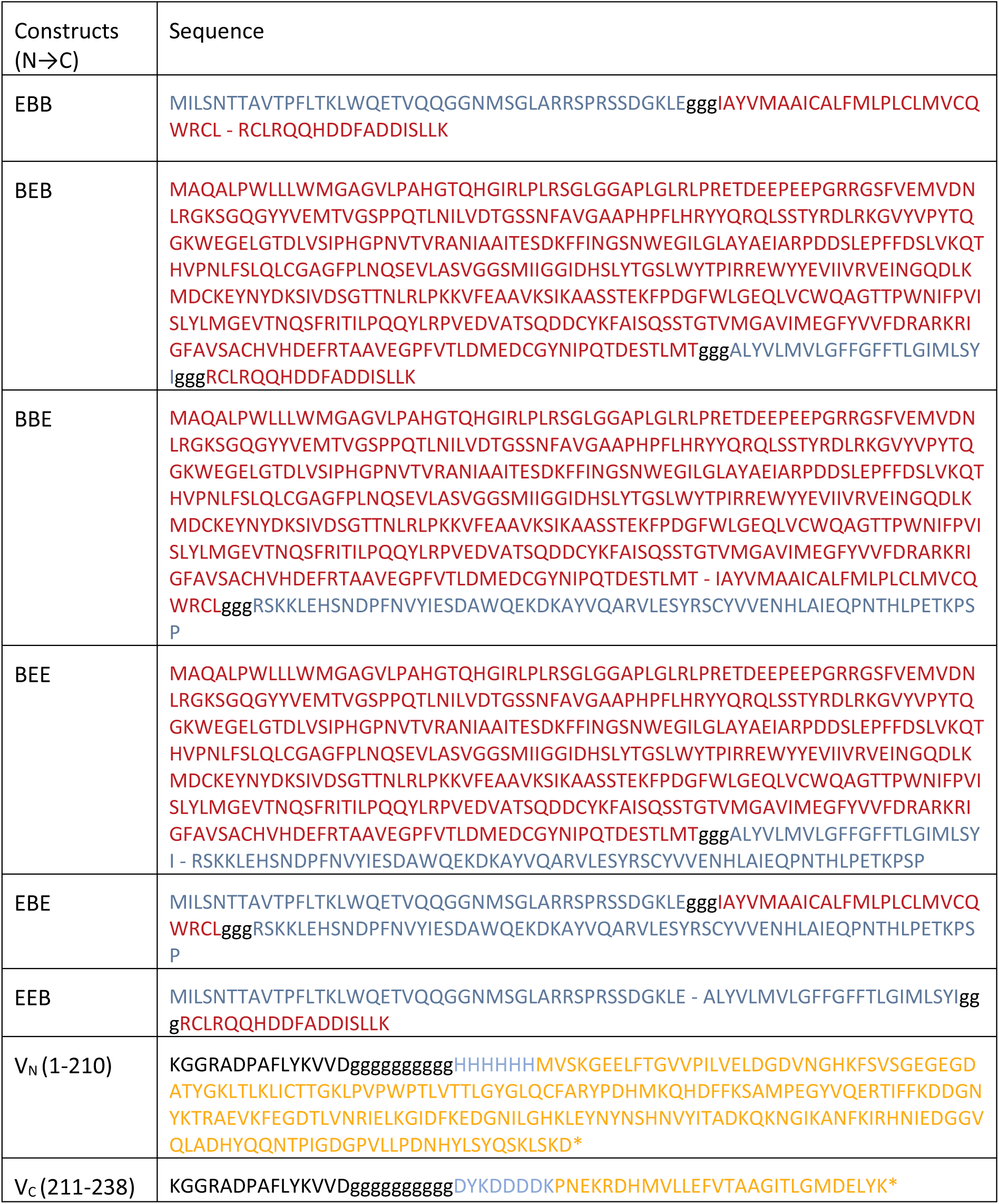
Amino acid sequences of BACE1–KCNE1 chimeric constructs and Venus BiFC fragments. Amino acid sequences of the BACE1–KCNE1 chimeras are shown, with extracellular, transmembrane, and intracellular domains indicated, together with the Venus N-terminal (V_N_) and C-terminal (V_C_) BiFC fragments. Colour coding: BACE1 (red), KCNE1 (blue), Venus (orange), vector backbone (black), His/FLAG tags (magenta). The intracellular C-terminus is indicated by an asterisk (*); glycine-rich linkers are shown in lowercase.

### Cell culture and transfection

HEK293T cells (ATCC® CRL-11268™) were maintained in DMEM (1 g/L glucose) supplemented with 10% fetal bovine serum (Biochrom) and 1% penicillin/streptomycin (Sigma-Aldrich) at 37 °C in a humidified atmosphere containing 5% CO₂.

For BiFC assays, 3 × 10⁵ cells were seeded onto poly-D-lysine–coated glass-bottom dishes (World Precision Instruments) 24 h prior to transfection. Cells were transfected with 200 ng of each BiFC construct using JetPEI (Polyplus Transfection) according to the manufacturer’s instructions.

For patch-clamp recordings, cells were plated on 35 mm tissue culture dishes (Falcon 353001, BD Biosciences) 24 h prior to transfection. Transfections were performed with JetPrime (Polyplus Transfection) using 150 ng KCNQ1, 150 ng KCNE1, 250 ng BACE1, and 75 ng EGFP (Clontech) per dish. After 24 h, cells were harvested with Accutase (Sigma-Aldrich), diluted 1:10 in fresh medium, and re-plated onto new 35 mm dishes for recordings.

For SiMPull assays, cells were seeded in 35 mm dishes and transfected the following day with 1.5 µg total DNA per dish using JetPrime. Samples were collected 24–36 h post-transfection.

For FRET assays, cells were plated on poly-D-lysine–coated (45 min) 1.5H borosilicate coverslips (VWR) and transfected 24 h later with 200 ng of each FRET construct using JetPEI (Polyplus Transfection).

### BiFC assay

Twenty-four hours after transfection, cells were overlaid with 1 mL HBS buffer and imaged on an LSM 780 confocal laser scanning microscope (Zeiss) equipped with a Plan-Apochromat 63×/NA 1.40 oil-immersion objective (Immersol 518F, Zeiss). Venus (YFP) fluorescence was excited at 514 nm, and emission was collected between 520–575 nm. To ensure direct comparability between samples, all imaging parameters were held constant: zoom factor 1.0, pixel dwell time 50.4 µs, line averaging 4×, master gain 1000, pinhole diameter 599 µm (12.5 AU), laser power 4%, and 16-bit acquisition at a resolution of 1024×1024 pixels.

### Electrophysiological Recordings

Potassium currents were recorded in whole-cell voltage-clamp configuration at room temperature (21 ± 1 °C) using a MultiClamp 700B amplifier and a Digidata 1320A interface controlled by Clampex 10.3 software (Molecular Devices). Pipettes were pulled from borosilicate glass capillaries with filament (BM150F-10, BioMedical Instruments) using a DMZ-Universal Puller (Zeitz Instruments) to a resistance of 2.5–3.5 MΩ in bath solution. Pipettes were filled on ice with intracellular solution using a 1 mL syringe (Soft-Ject, Henke-Sass, Wolf) connected to a nylon filter (0.2 µm pore, Nalgene, Thermo Fisher Scientific) and a Microfil (34 G, 67 mm, World Precision Instruments). The intracellular solution contained (in mM): 135 K-gluconate, 10 KCl, 4 NaCl, 5 HEPES, 5 EGTA, 2 Na₂-ATP, and 0.3 Na₃-GTP (all Sigma-Aldrich), adjusted to pH 7.25 with KOH (Carl Roth). The extracellular solution contained (in mM): 145 NaCl (Carl Roth), 4 KCl (Merck), 2 CaCl₂·2H₂O (Sigma-Aldrich), 2 MgCl₂·6H₂O (Merck), 10 D-glucose·H₂O (Merck), and 10 HEPES (Sigma-Aldrich), adjusted to pH 7.4 with NaOH (Merck). Access resistance was ≤ 6 MΩ before series resistance compensation (≥75%). Recordings were sampled at 20 kHz and filtered at 6 kHz. No liquid junction potential correction was applied.

### Single Molecule Pull-down (SiMPull) assay

SiMPull assays were performed as described previously (Jain et al., 2011), with minor modifications. HEK293T cells were co-transfected with an HA-tagged bait protein (HA-hBACE1 or HA-hKCNQ1) and a hBACE1 prey protein, either with or without an α-bungarotoxin labeling site (hBACE1-BBS or native hBACE1). Twenty-four hours post-transfection, cells were incubated with 1 ng/µL α-bungarotoxin-ATTO Fluor-647 (Alomone) for hBACE1-BBS labeling or with 0.1 µM Alexa568-C3 for native hBACE1 (Stockinger et al., 2024) for 20 min at room temperature. Cells were washed twice with culture medium (20 min each) and lysed in buffer containing (in mM): 150 NaCl, 10 Tris (pH 7.5), 1 EDTA, 1.5% IGEPAL (Sigma), and a protease inhibitor cocktail (Thermo Scientific). To remove free fluorophores, lysates were purified using PD MiniTrap G-10 desalting columns (Cytiva) equilibrated in T50 buffer (in mM: 50 NaCl, 10 Tris, 20 EDTA; 0.1 mg/mL BSA; pH 7.5). Purified lysates were applied to PEG-passivated coverslips (99% PEG, 1% biotin-PEG) coated sequentially with neutravidin (1.4 mg/mL, Pierce) and biotinylated anti-HA antibody (15 nM; Abcam, #ab26228). After extensive washing with T50 buffer to minimize nonspecific binding, single-molecule complexes were visualized by total internal reflection fluorescence microscopy using a 100× objective (Olympus) and using 642 nm and 561 nm excitation, as appropriate. Movies (13 × 13 µm², 250 frames, frame rates of 10–30 Hz) were analysed with Fiji software (NIH). Only spots that photobleached completely during illumination were included. Data were obtained from multiple independent experiments, with a minimum of 15 movies per condition and at least 50 analysed traces per movie. Representative images are shown; quantification includes all datasets/experiments.

### FRET assay

HEK293T cells were fixed 48 h post-transfection with PBS containing 4% paraformaldehyde (C. Roth), washed repeatedly with PBS, and stained with 300 nM Alexa568-C3 and 100 nM Alexa488-C3 (Stockinger et al., 2024) for 15 min at room temperature. Coverslips were mounted in Roti-Mount FluorCare (C. Roth), sealed with Twinseal (picodent GmbH), and stored at 4 °C for a maximum of 24 h before imaging. FRET measurements were performed on an LSM 780 confocal microscope (Zeiss) equipped with a Plan-Apochromat 63×/NA 1.40 oil-immersion objective, an argon laser LGN 3001 (LASOS), and a DPSS 561-10 laser system. Fluorescence emission was recorded in lambda mode and extracted from two custom-defined spectral windows (516–524 nm and 595–603 nm). All acquisition parameters were identical across experiments. For each cell, mean fluorescence intensities were quantified before and after acceptor photobleaching. Background signals were determined from a cell-free region of interest (ROI) and subtracted accordingly. FRET efficiencies were calculated and corrected for Alexa488/Alexa568 stoichiometry as described previously (Stockinger et al., 2024).

### Data analysis and statistics

Data analysis and visualization were performed using OriginPro 2015G 64-bit (OriginLab Corporation). Unless otherwise stated, values are presented as mean ± standard error of the mean (SEM). For non-normally distributed data, the geometric mean was used. In these cases, the arithmetic mean ± SEM was calculated from log-transformed data, and individual values were subsequently back-transformed for graphical representation. Statistical tests and confidence intervals are specified in the corresponding figure legends.

### Use of AI tools

OpenAI’s ChatGPT (GPT-5; accessed January 2026) was used solely for language editing—to improve clarity, grammar, and readability. All substantive content, analyses, and interpretations were independently conceived and verified by the authors, who accept full responsibility for the manuscript.

## Results

### The BiFC assay confirms physical interaction of KCNE1 and BACE1 with KCNQ1

In our previous work, we showed that the β-secretase BACE1 interacts directly with KCNQ1 and neuronal KCNQ channels in a non-proteolytic manner (Agsten et al., 2015; Hessler et al., 2015). Here, we elaborated on these findings to gain deeper insight into the interplay between KCNQ1, its canonical subunit KCNE1, and BACE1 at the molecular level. To probe direct physical interaction, we employed a bimolecular fluorescence complementation (BiFC) assay using yellow fluorescent protein (YFP/Venus) (Ohashi et al., 2012). Split Venus fragments, either V_N_ (1-210) or V_C_ (211-238), were attached to the C-termini of the proteins examined.

First, we validated the BiFC assay by confirming the homotetrameric assembly of KCNQ1 α-subunits (Pusch et al., 2000) (Fig. 1A). As a control, we used a BiFC-tagged epithelial sodium channel subunit ENaC1α. To our knowledge, no direct physical interactions between ENaC and KCNQ1 have been reported. However, ENaC1α forms oligomeric complexes (Cheng et al., 1998), producing a relatively weak BiFC signal (Fig. 1B). These experiments confirmed that KCNQ1 and ENaC1α constructs carrying BiFC tags were expressed and capable of forming homomeric assemblies. Co-expression of ENaC1α with KCNQ1 in both BiFC combinations yielded only very weak fluorescence signals (Fig. 1C-D). The low-level signal most likely resulted from random encounters in densely packed cellular compartments, importantly, no signal was detected at the plasma membrane. By contrast, the homotetrameric assembly of KCNQ1 α-subunits yielded a strong signal above background (Fig. 1A, E-F).

**Figure 1.**
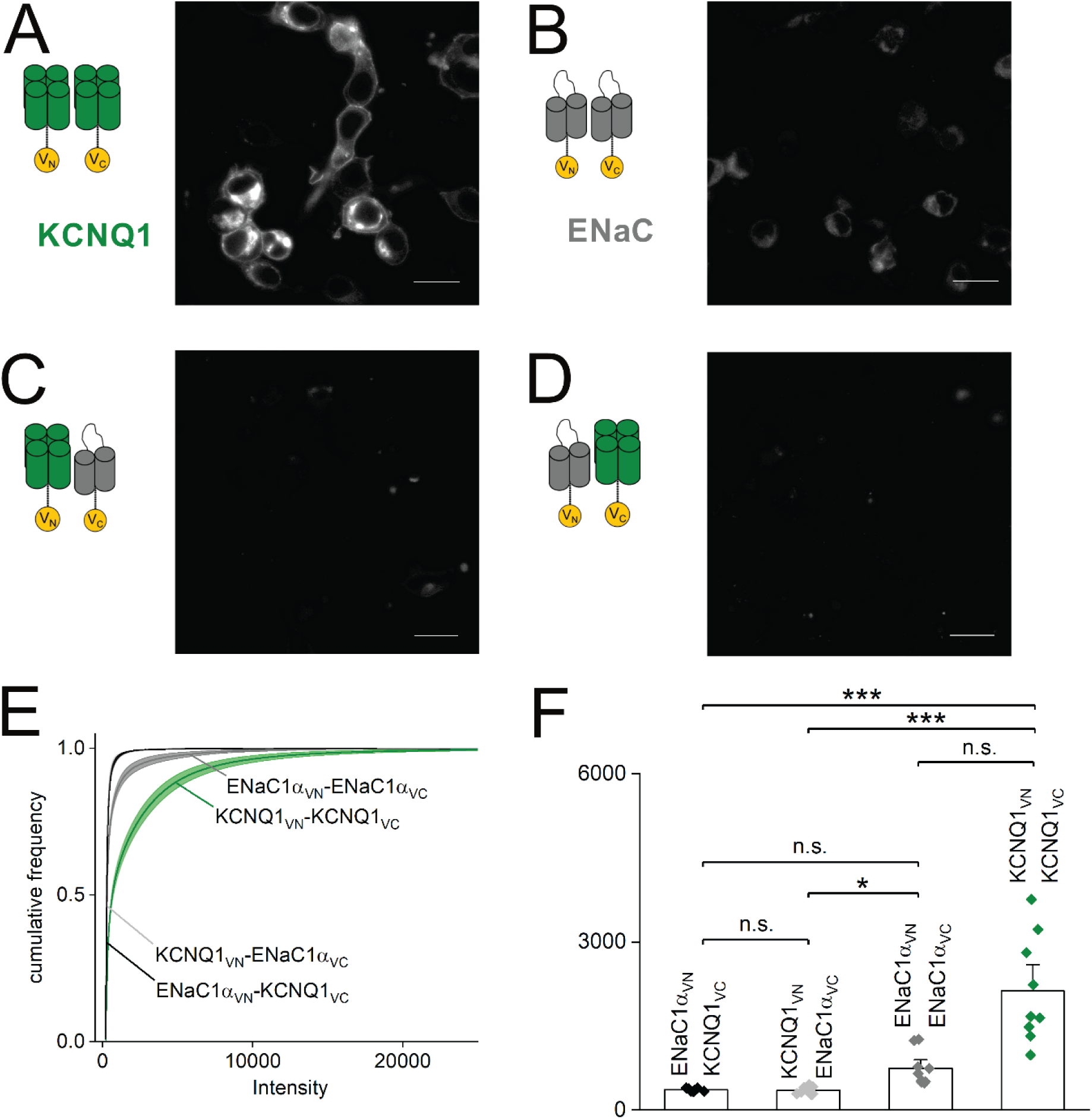
Validation of the bimolecular fluorescence complementation assay (BiFC) – confirming homomeric assembly of KCNQ1. **(A-D)** HEK293T cells were transfected with the constructs indicated on the left side of the images acquired by laser scanning. The ENaC1α construct served as control. For each experiment, one construct was tagged at the C-terminus with either the long split Venus fragment V_N_ or the shorter V_C_ construct. Assembly of V_N_ and V_C_ constructs resulted in a fluorescence signal. **(E)** After removing background fluorescence, intensity distributions were generated and depicted as cumulative probabilities. **(F)** Mean intensities were used for comparison of construct interaction. Scale bars represent 20 µm. n = 10 images each from two independent transfections. Kruskal–Wallis, with Dunn’s test for post hoc pairwise comparisons. *p < 0.05, ***p < 0.001.

From the recorded images, we generated background-subtracted cumulative intensity distributions (Fig. 1E). Although these curves capture the full spectrum of interaction strengths, they can be difficult to interpret directly. Therefore, we instead compared mean fluorescence intensities (Fig. 1F). ENaC1α co-expressed with KCNQ1 served as our negative control, and any mean intensity significantly different from this control served as evidence of a specific interaction.

Next, we examined the physical interaction of KCNQ1 with its canonical subunit KCNE1 (Fig. 2A-B, E-F) and with BACE1 (Fig. 2C-D, G-H). When co-expressed with KCNE1-V_C_ or BACE1-V_C_, the KCNQ1-V_N_ construct, which carries the longer YFP fragment, displayed a fluorescence signal barely above ENaC1α control levels (Fig. 2A-B), suggesting a possible steric hindrance in this orientation. We further demonstrated that the KCNQ1 and KCNE1 tagged constructs were fully functional, as they were expressed at the plasma membrane and gave rise to typical KCNQ1 currents (Fig. 2I) and *I_Ks_* (Fig. 2J-K), respectively.

**Figure 2.**
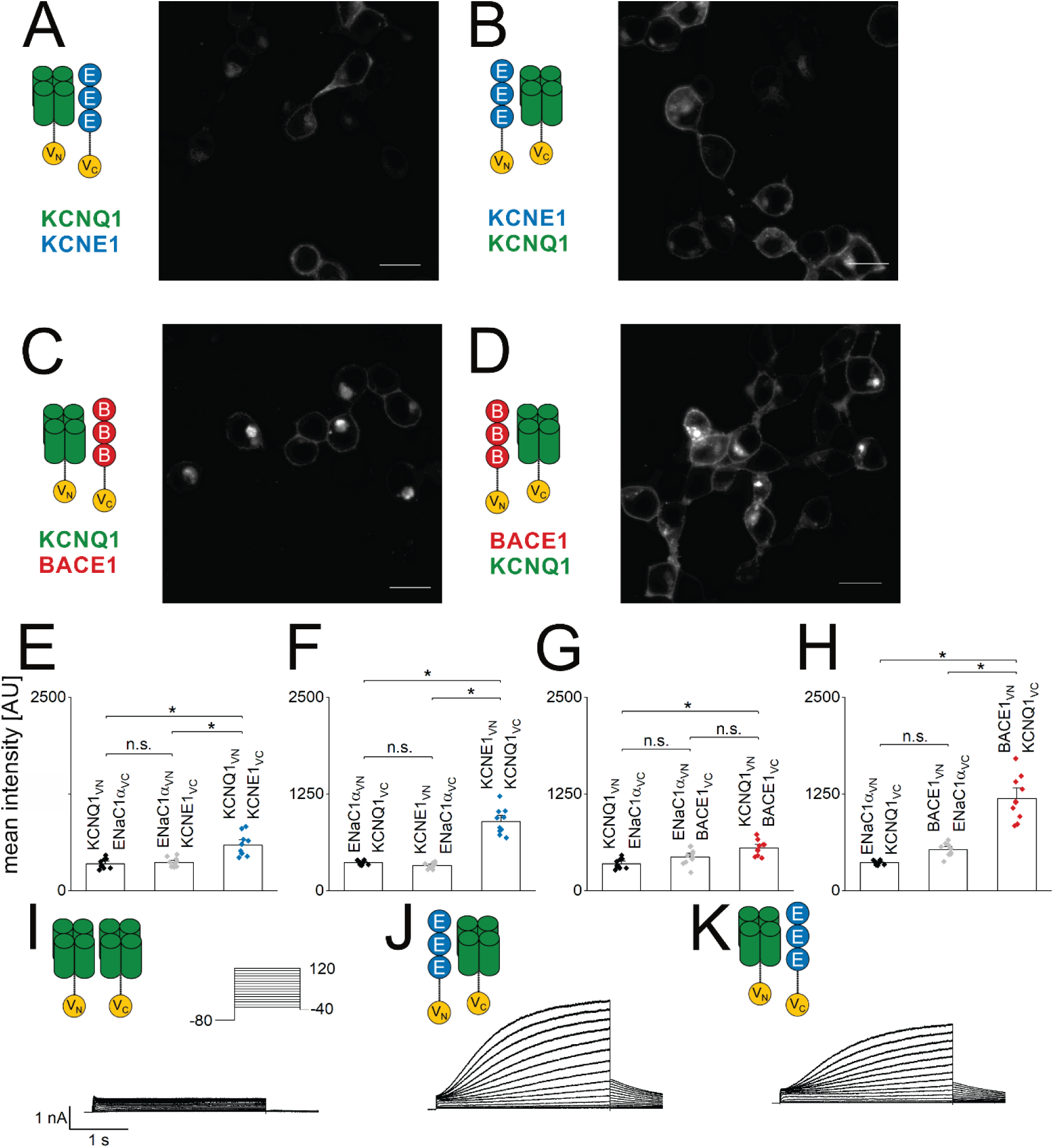
The BiFC assay reveals direct interaction of KCNQ1 with KCNE1 and with BACE1. **(A-D)** HEK293T cells were transfected with the constructs indicated on the left side of the laser-scanning images tagged with either the long (V_N_) or short (V_C_) BiFC construct. Fluorescence emission after assembly of the BiFC constructs was recorded by confocal laser scanning. Scale bars represent 20 µm. **(E-H)** Mean intensities were compared to control experiments with ENaC1α. n = 10 images out of two independent transfections. Kruskal–Wallis with Dunn’s test for post hoc pairwise comparisons. *p < 0.05. **(I-K)** Patch-clamp recordings revealed the typical KCNQ1 and KCNQ1/E1 (I_Ks_) current phenotype, respectively, indicating that neither assembly nor trafficking of the channel complexes was grossly impaired by the BiFC tags. n = 3 recordings for each combination.

### Physical and functional interactions of KCNE1-BACE1 chimeric constructs with KCNQ1

In the following experiments, we employed the BiFC assay to pinpoint the regions of KCNE1 and BACE1 mediating their physical association with the KCNQ1 pore-forming subunit. We engineered six chimeric proteins by exchanging the N- and C-terminal interfaces flanking the transmembrane helices of KCNE1 and BACE1 (see Methods; Fig. 3). Each chimera carried one half of the split Venus protein, while full-length KCNQ1 bore the complementary fragment on its C-terminus. As a specificity control for comparison, we measured the BiFC signal between KCNQ1 and the unrelated ENaC1α subunit.

**Figure 3.**
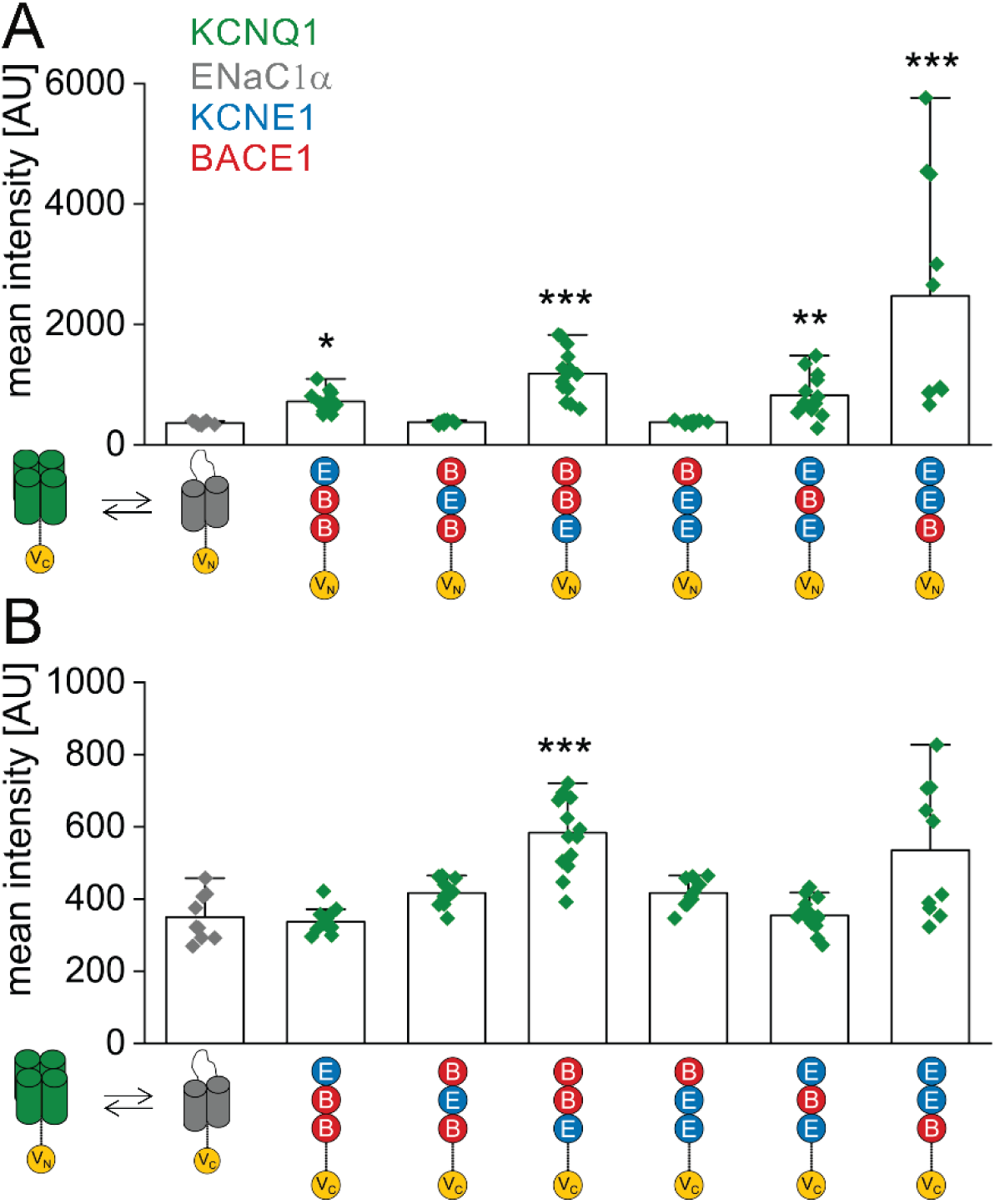
Elucidating the KCNE1–BACE1 interaction domains with KCNQ1 using the BiFC assay. Six chimeric constructs were designed, each comprising the extracellular (ECD), transmembrane (TMD), and intracellular (ICD) domain of KCNE1 or BACE1, fused C-terminally to V_N_ or V_C_. HEK293T cells were co-transfected with each chimera and full-length KCNQ1 carrying the complementary Venus fragment. ENaC1α served as a negative control. In **(A)**, KCNQ1 carried V_C_; in **(B)**, KCNQ1 carried V_N_. Fluorescence intensities were measured by laser-scanning confocal microscopy; data are mean ± SEM; n = 10 from two independent transfections per pairing. Statistical significance was assessed by Kruskal–Wallis test with Dunn’s post hoc multiple-comparisons test. *p < 0.05, **p < 0.01, ***p < 0.001.

Consistent with our previous observations, KCNQ1 tagged with the shorter VC fragment produced higher overall fluorescence intensities (Fig. 3A). Among the six constructs tested, the chimeras with either BACE1 or KCNE1 in both the extracellular and transmembrane domains (BBE and EEB) showed the highest fluorescence signal (Fig. 3A-B), although the signal for EEB did not reach statistical significance in panel B. Chimeras with KCNE1 in the extracellular domain and BACE1 in the transmembrane domain (EBB and EBE) also yielded significant BiFC signals when fused to the VN tag (Fig. 3A). In contrast, constructs combining the extracellular domain of BACE1 with the transmembrane domain of KCNE1 (BEB and BEE) failed to produce fluorescence above background levels (Fig. 3A-B).

These data indicate that most BACE1-KCNE1 chimeras can physically interact with KCNQ1, reflecting an appreciable degree of domain compatibility, whereas pairing BACE1’s extracellular region with KCNE1’s transmembrane segment precludes this interaction.

To further dissect the functional interplay of our chimeric constructs, we performed electrophysiological recordings of KCNQ1 currents and organized the results into two distinct phenotypic groups. In Figure 4, we illustrate currents exhibiting a KCNQ1-like phenotype by comparing recordings from KCNQ1 alone with those from KCNQ1 co-expressed with wild-type BACE1 (Fig. 4A-B). By contrast, Figure 5 depicts an *I_Ks_*-like phenotype, showing currents from KCNQ1 co-expressed with KCNE1 alone as well as those from KCNQ1 co-expressed with both KCNE1 and wild-type BACE1 for reference (Fig. 5A-B).

**Figure 4.**
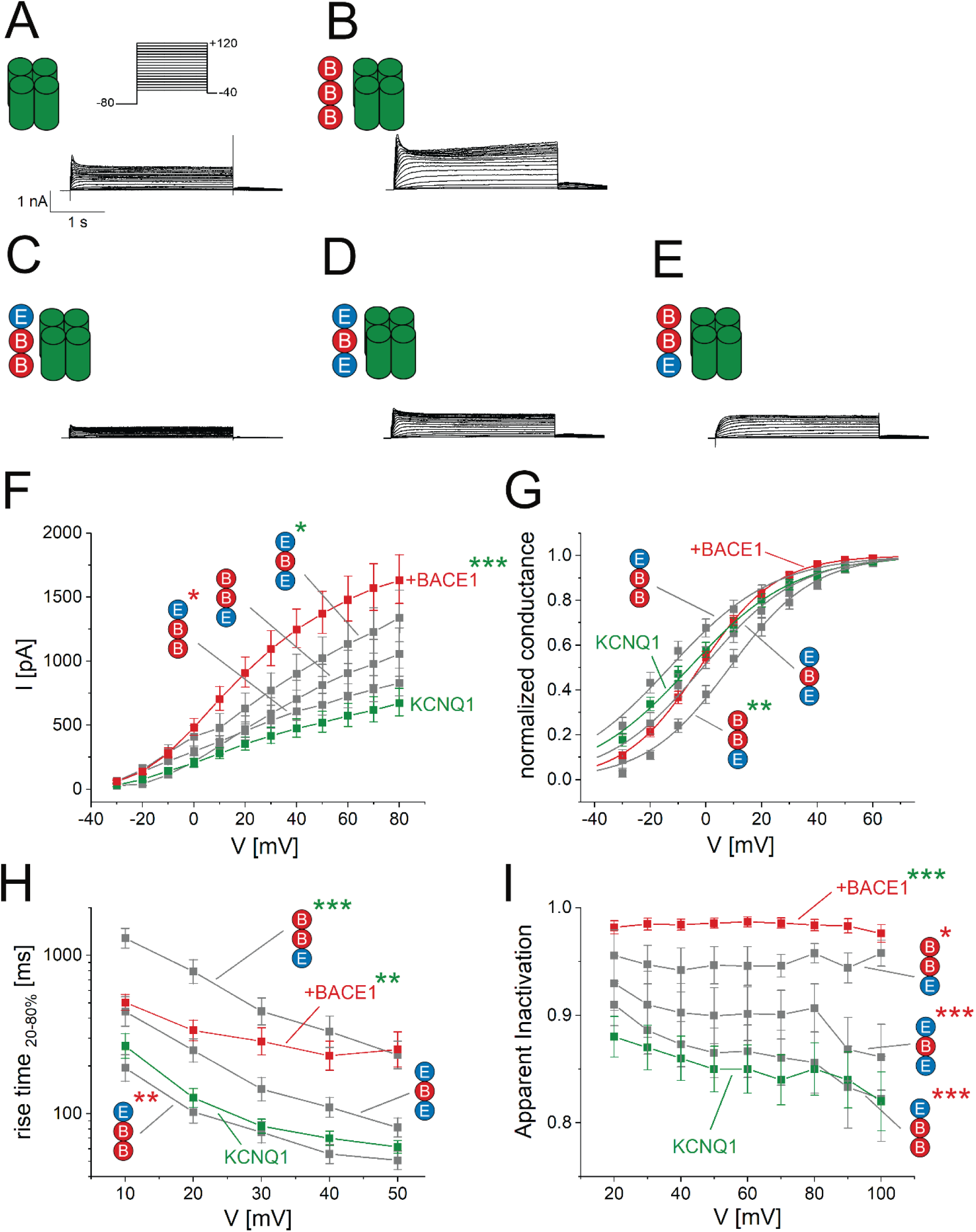
Electrophysiological characterisation of KCNQ1 currents together with KCNE1-BACE1 chimeric constructs containing the BACE1 transmembrane domain revealed KCNQ1 currents without *I_Ks_*-like characteristics. Chimeric constructs were designed to contain the complete extracellular (ECD) and intracellular (ICD) domain of either KCNE1 or BACE1, respectively, and the transmembrane domain of BACE1. HEK293T cells were transfected with untagged chimeric constructs in addition to KCNQ1. (A-C) Typical patch-clamp recordings with step protocols are depicted as indicated. From these recordings the current-voltage relation (D), the normalized conductance (E), the current rise time (F), and the fraction of inactivation (G) were quantified. Groups were compared point-wise against KCNQ1 and KCNQ1 + BACE1 using a Kruskal-Wallis test, followed by Dunn’s post hoc pairwise comparisons with p-values adjusted for multiple testing. Sample sizes (maximum per group; smaller at specific points due to dropouts): KCNQ1, n=19; KCNQ1 + BACE1, n=14; BBE, n=13; EBB, n=16; EBE, n=16. Asterisks denote adjusted pairwise p-values and are shown only when at least two data points meet the threshold: *p < 0.05, **p < 0.01, ***p < 0.001.

**Figure 5.**
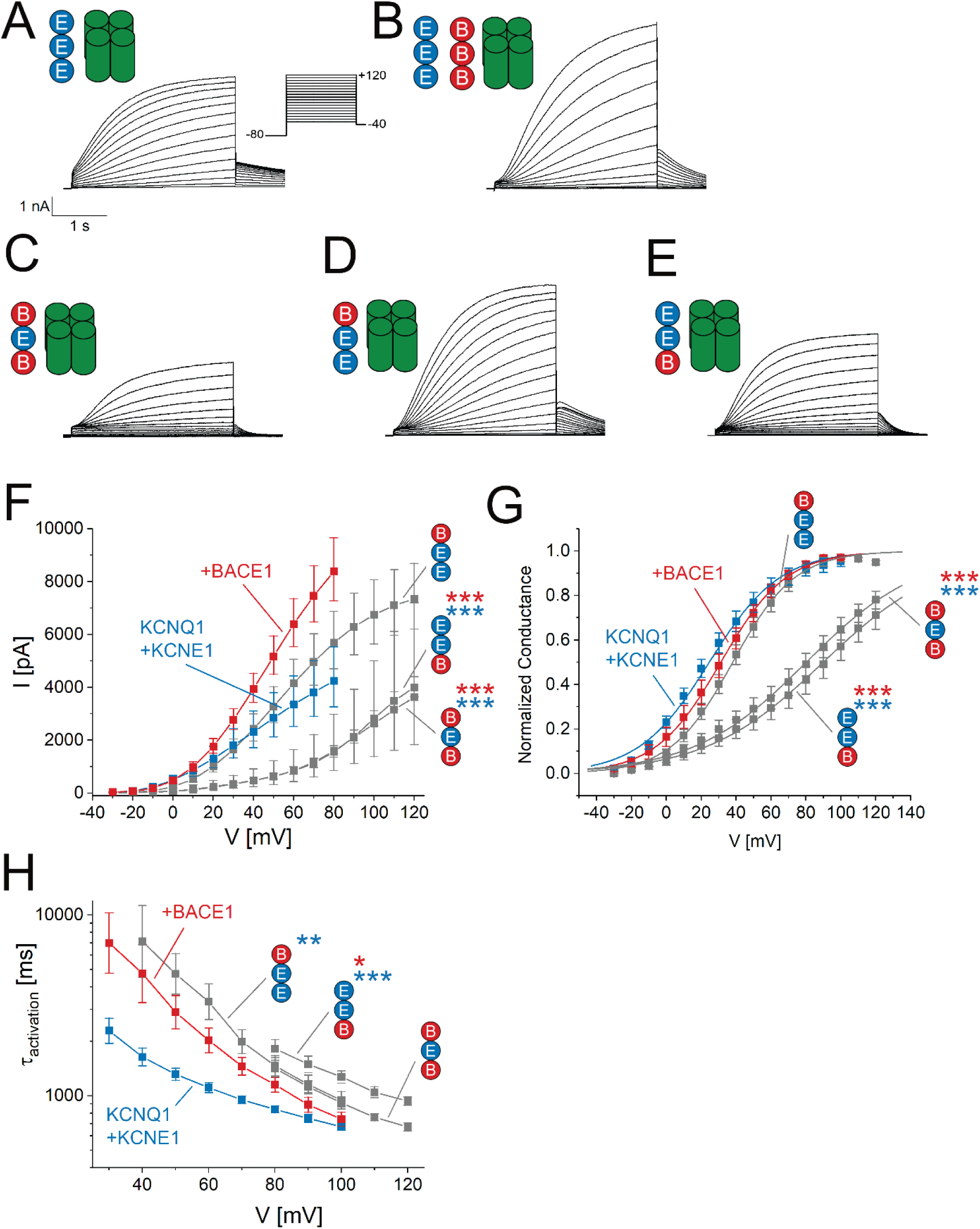
Electrophysiological characterisation of KCNQ1 currents together with KCNE1-BACE1 chimeric constructs containing the KCNE1 transmembrane domain revealed *I_Ks_*-like currents. Chimeric constructs were designed to contain the complete extracellular (ECD) and intracellular (ICD) domain of either KCNE1 or BACE1, respectively, and the transmembrane domain of KCNE1. HEK293T cells were transfected with untagged chimeric constructs in addition to KCNQ1. (A-C) Typical patch-clamp recordings with step protocols are depicted as indicated. From these recordings the current-voltage relation (D), the normalized conductance (E), and the current activation time constant (F) were quantified. Groups were compared point-wise against KCNQ1 + KCNE1 and KCNQ1 + KCNE1 + BACE1 using a Kruskal-Wallis test, followed by Dunn’s post hoc pairwise comparisons with p-values adjusted for multiple testing. Sample sizes (maximum per group; smaller at specific points due to dropouts): KCNQ1 + KCNE1, n=18; KCNQ1 + KCNE1 + BACE1, n=18; BEE, n=19; EEB, n=26; BEB, n=25. Asterisks denote adjusted pairwise p-values and are shown only when at least two data points meet the threshold: *p < 0.05, **p < 0.01, ***p < 0.001.

Each wild-type accessory subunit uniquely modulated KCNQ1 current amplitude, and co-transfection of both subunits produced an additive effect on current traces. KCNE1 shifted the voltage dependence of activation toward more depolarized potentials (compare Figs. 4G and 5G), whereas BACE1 markedly slowed activation kinetics (Figs. 4H, significant and 5H, not significant) without substantially altering voltage-dependent activation (Figs. 4G and 5G). Consistently, BACE1 most strongly affected KCNQ1 gating by slowing activation (Agsten et al., 2015). Notably, unlike this previous report, we did not observe notable inactivation in the presence of BACE1. Apparent inactivation of KCNQ1 is known to be variable (reviewed in Wang et al., 2020) and reflects the balance between activation and inactivation kinetics. In our recordings of KCNQ1, inclusion of ATP in the intracellular solution likely slowed activation sufficiently that apparent inactivation was no longer evident.

Building on these controls, we then evaluated recordings from our panel of chimeric subunits. Five constructs produced distinct alterations in KCNQ1 gating, whereas the chimera containing the KCNE1 extracellular domain fused to BACE1’s transmembrane and intracellular regions (EBB) had no significant effect on channel behaviour (Fig. 4C, 4F-I). The EBE chimera induced only subtle gating changes (Fig. 4D, 4F-I). In contrast, the BBE chimera, housing BACE1’s extracellular domain in place of KCNE1, faithfully reproduced the slowed activation kinetics and lack of inactivation characteristic of wild-type BACE1 (Fig. 4E, 4H, 4I – not significant).

Consistent with a previous study (Melman et al., 2001), the transmembrane domain was necessary and sufficient to confer a KCNE1-like phenotype (Fig. 5C-E). Additionally, chimeras incorporating the KCNE1 transmembrane segment (BEB, BEE, EEB) displayed the characteristic triad of depolarizing shift in voltage-dependence (Fig. 5G), absence of inactivation, and slowed activation kinetics (Fig. 5H). Furthermore, inclusion of the KCNE1 intracellular domain was required to fully recapitulate wild-type KCNE1 currents: constructs lacking this domain (BEB, EEB) activated at excessively positive potentials (Fig. 5G). Remarkably, the BEE chimera, combining the BACE1 extracellular region with both the KCNE1 transmembrane and intracellular modules, emerged as a “super-mutant,” exhibiting additive effects of both accessory subunits on gating, namely characteristic *I_Ks_* current with slowed activation (Fig. 5H).

Together, these results indicate that BACE1 affects KCNQ1 gating primarily by its large extracellular domain, whereas KCNE1’s influence depends on its transmembrane segment, with its intracellular domain fine-tuning the overall phenotype.

### BACE1 stoichiometry in KCNQ1 complexes and relationship to KCNE1

The number of KCNE1 subunits incorporated into a KCNQ1 channel complex is a long-standing matter of debate. Recent studies argue in favour of a variable stoichiometry of up to four KCNE1 subunits recruited into a homotetrameric KCNQ1 complex (reviewed in Wang et al., 2020). Our previous electrophysiological recordings (Agsten et al., 2015) suggested that KCNE1 and BACE1 can modulate KCNQ1 simultaneously. Because BACE1 also binds directly to KCNQ1 (Fig. 2C-D, G-H), the question of binding stoichiometry becomes more complex: KCNE1 and BACE1 may compete for overlapping or adjacent sites on the channel, with potential consequences for both subunit incorporation and overall channel function.

To investigate the association between BACE1 and KCNQ1 channel complexes, we employed the single-molecule pull-down (SiMPull) assay (Jain et al., 2011; Levitz et al., 2016; Royal et al., 2019). In this approach, antibody-mediated immobilization of individual protein complexes on passivated coverslips enabled us to determine stoichiometry by counting discrete photobleaching steps of fluorescent labels.

In a first experiment, an HA-BACE1 fusion protein was co-transfected with a BACE1 protein containing an α-bungarotoxin binding site (BACE1-BBS) that was labelled with an ATTO Fluor-647 bungarotoxin conjugate (Fig. 6A). Following HA-affinity capture on passivated coverslips, we detected only minimal fluorescence in control experiments using lysates from non-transfected cells or from cells expressing BACE1-BBS alone (Fig. 6B-C). By contrast, co-expression of HA-BACE1 and BACE1-BBS produced numerous fluorescent puncta (Fig. 6D), consistent with pull-down of BACE1-BBS with HA-BACE1 and thus confirming BACE1-BACE1 association (Liebsch et al., 2017; Stockinger et al., 2024). Finally, co-transfection of HA-KCNQ1 with BACE1-BBS yielded a similarly robust fluorescence signal (Fig. 6E), demonstrating a direct physical interaction between BACE1 and the KCNQ1 channel complex with the SiMPull assay.

**Figure 6.**
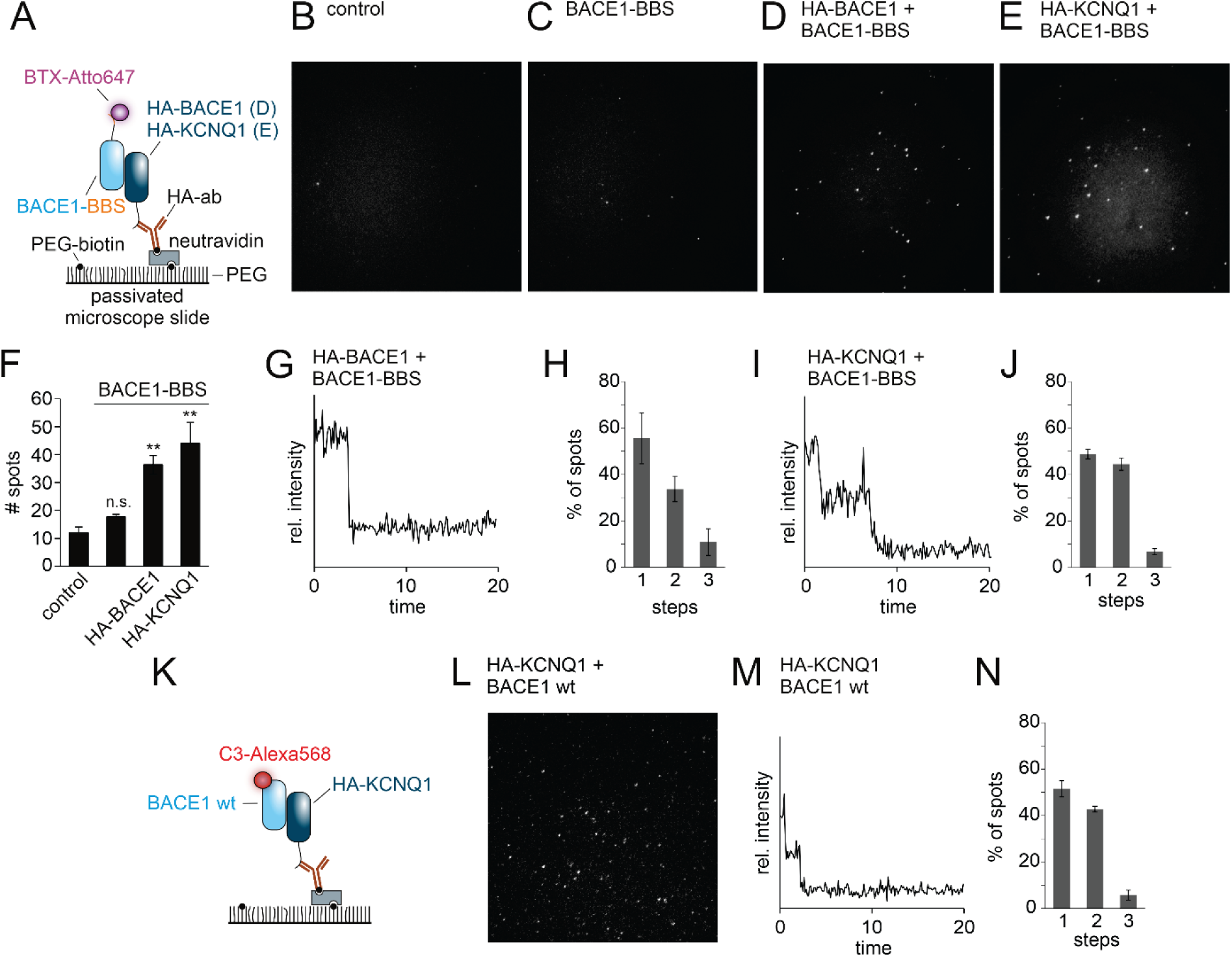
Single-molecule pull-down (SiMPull) of BACE1 and KCNQ1. **(A)** Schematic of the SiMPull assay. HEK293T cells were co-transfected with BACE1 carrying a bungarotoxin binding site (BACE1-BBS) and either HA-BACE1 or HA-KCNQ1. Cell lysates were applied to PEG-passivated coverslips coated with biotinylated anti-HA antibodies to capture HA-tagged proteins and their interaction partners. BACE1-BBS was detected by labeling with ATTO647-conjugated α-bungarotoxin (BTX-ATTO647). **(B)** TIRF image of control (non-transfected cells). **(C)** TIRF image of cells expressing only BACE1-BBS, showing low background labeling. **(D, E)** TIRF images of specific pull-down events, where BACE1-BBS was co-captured with HA-BACE1 (D) or HA-KCNQ1 (E). **(F)** Quantification of spot counts from condition shown in (B-E). **(G)** Example fluorescence trace with a single photobleaching step, indicating one ATTO647 fluorophore on BACE1-BBS co-captured with HA-BACE1. **(H)** Photobleaching step distribution for BACE1-BBS with HA-BACE1. n = 6 independent experiments. **(I)** Example trace with two photobleaching steps, indicating two BACE1-BBS co-captured with HA-KCNQ1. n = 3 independent experiments. **(J)** Photobleaching step distribution for BACE1-BBS with HA-KCNQ1. **(K)** Schematic of a complementary assay using wild-type BACE1 (BACE1 wt) co-expressed with HA-KCNQ1. Here, BACE1 wt was affinity labeled with an Alexa568-conjugated BACE1 inhibitor (C3-Alexa568) and captured via HA-KCNQ1. **(L)** Representative TIRF image showing pull-down of C3-Alexa568–labeled BACE1 wt with HA-KCNQ1. **(M)** Example trace with two photobleaching steps of labeled BACE1 wt. **(N)** Photobleaching step distribution for BACE1 wt with HA-KCNQ1. n = 6 independent experiments. Data were analysed using one-way ANOVA, with Tukey’s test for post hoc pairwise comparisons. **p < 0.01.

Bleaching-step analysis of individual fluorescent puncta revealed the subunit composition of the captured complexes. In the HA-BACE1 / BACE1-BBS SiMPull assay (Fig. 6G-H), ≈56% of spots bleached in a single step, while ≈34% showed two steps and ≈11% three steps (Fig. 6H). These findings add weight to earlier reports of dimeric and trimeric BACE1 assemblies (Liebsch et al., 2017; Schmechel et al., 2004; Stockinger et al., 2024; Westmeyer et al., 2004) and underscore the functionality of the assay.

When probing the HA-KCNQ1 / BACE1-BBS interaction, an almost equal number of spots bleached in one step (≈49%) and in two steps (≈44%), with only ≈7% displaying three steps (Figs. 6I-J). To corroborate this stoichiometry, we repeated SiMPull using wild-type BACE1 co-expressed with HA-KCNQ1, detecting BACE1 via the novel inhibitor-based probe C3-Alexa568 (Fig. 6K-L) (Stockinger et al., 2024). The resulting bleaching-step distribution, ≈52% one-step, ≈43% two-step, and ≈6% three-step events (Figs. 6M-N), mirrored the above observations. Given a labelling efficiency of about 0.6 for C3-Alexa568 (Stockinger et al., 2024), and assuming a similar efficiency for ATTO Fluor-647 bungarotoxin because of a similar bleaching step distribution, we conclude that predominantly two BACE1 molecules bind to each KCNQ1 complex. The small fraction of three-step events could reflect occasional co-detection of two diffraction-limited, independently captured complexes within a single spot rather than a true 3:1 stoichiometry.

Altogether, these data lend strong support to the view that BACE1 manifests as an oligomeric complex of two or three molecules (Stockinger et al., 2024). Moreover, our findings demonstrate a predominantly two-to-one binding stoichiometry of BACE1 with the KCNQ1 channel.

In a complementary set of experiments, we assessed BACE1 recruitment to the KCNQ1 channel complex using the BiFC assay as outlined above. HEK293T cells were transfected with BACE1 V_N_ and BACE1 V_C_ constructs either alone or together with untagged KCNQ1. Pairing the two BACE1 constructs generated a robust plasma membrane BiFC signal indicating BACE1 homomeric assembly (Fig. 7A). KCNQ1 co-transfection did not alter the subcellular distribution of this signal but induced an overall reduction in BiFC intensity (Fig. 7B-C). We take this observation as further evidence for a direct physical interaction between BACE1 and KCNQ1, with the novel implication that the presence of KCNQ1 weakens the homomeric assembly of BACE1.

**Figure 7.**
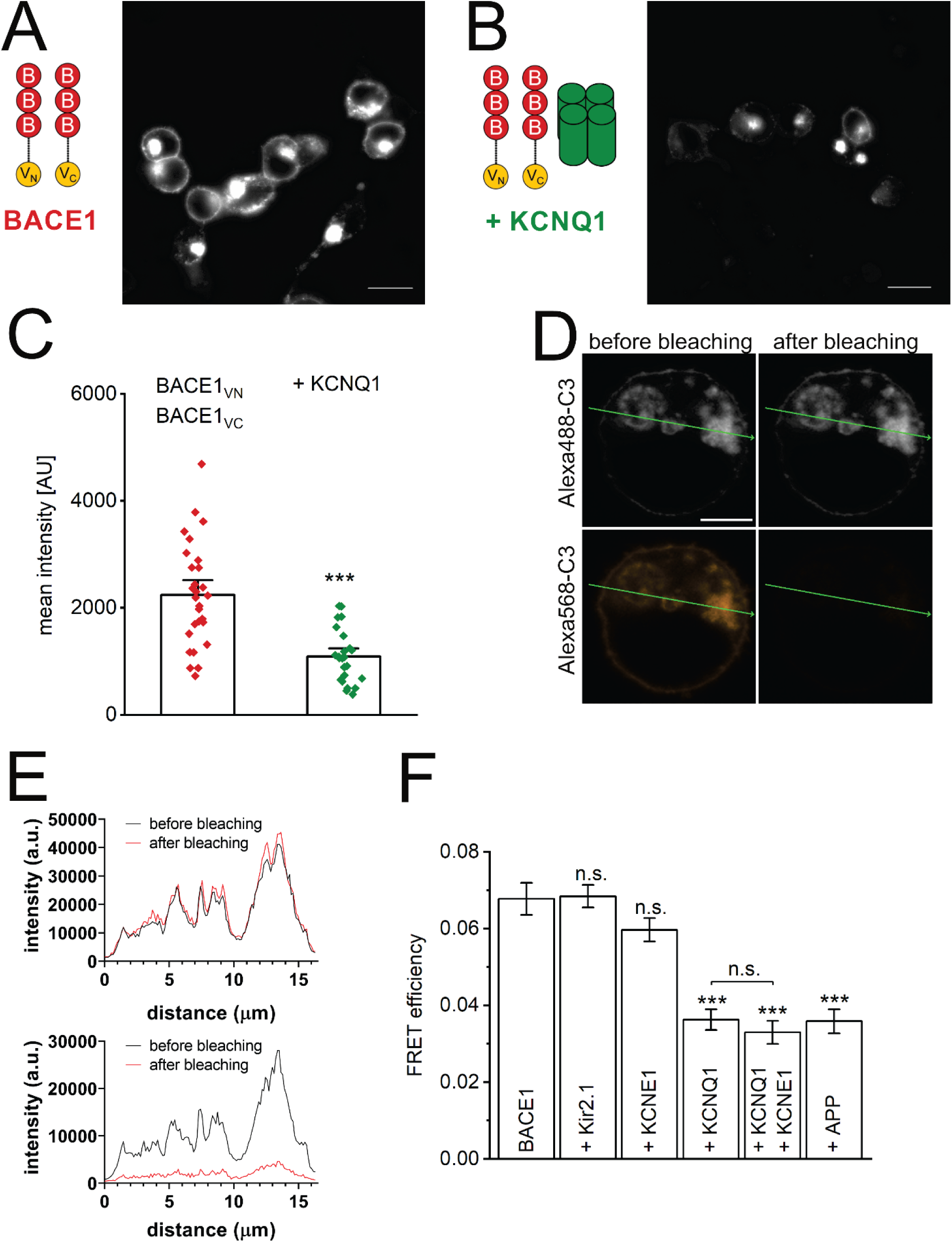
Co-expression of KCNQ1 reduces BACE1 homomeric assembly. **(A, B)** Laser scanning images show representative dimerization of BACE1 BiFC constructs transfected in HEK293T cells. (B) Additionally, KCNQ1 was co-transfected. **(C)** Images as depicted in (A,B) were quantified using mean BiFC intensities of BACE1 dimers. n = 28 (BACE1); n = 24 (+KCNQ1) cells from eight independent transfections, Mann-Whitney U test, ***p < 0.001. **(D-F)** BACE1 homomeric assembly in the absence or presence of KCNQ1 was additionally assessed by acceptor photobleaching FRET (Förster resonance energy transfer). HEK293T cells were transfected with BACE1 wild-type constructs and with and without KCNQ1. BACE1 was labeled with the small molecule BACE1 inhibitor C3 coupled to either Alexa488 serving as donor or Alexa568 serving as acceptor in a ratio of 1:3. **(D)** Typical fluorescence images before (left panel) and after (right panel) photobleaching are shown with the Alexa488 channel in the top panel and the Alexa568 channel in the panel below, respectively. The fluorescence profile along the green arrow is depicted in **(E)** The Alexa488 channel shown in the top graph indicates an intensity gain after bleaching. The Alexa568 channel in the graph below indicates almost complete bleaching of the acceptor. **(F)** Calculated FRET efficiency for BACE1 multimers is visualized in the graph and additionally with co-transfection of potassium channel Kir2.1, the subunit KCNE1, KCNQ1 + KCNE1 and amyloid precursor protein (APP). n = 34 (BACE1); n = 24 (+ Kir2.1); n = 40 (+ KCNE1); n = 17 (+ KCNQ1); n = 23 (+ KCNQ1 + KCNE1); n = 23 (+ APP) cells from two independent transfections. One-way ANOVA, ***p < 0.001.

To validate the latter finding independently, we employed acceptor photobleaching Förster resonance energy transfer (FRET) with the BACE1 inhibitor-derived probes C3-Alexa488 and C3-Alexa568 at a 1:3 donor-to-acceptor ratio. Cells expressing wild-type BACE1 were imaged before and after bleaching the Alexa568 acceptor (Fig. 7D-E). The resulting increase in Alexa488 donor fluorescence revealed donor dequenching and enabled calculation of corrected FRET efficiencies (Fig. 7F). As a control we used the Kir2.1 potassium channel. Kir2.1 co-expression had no effect on the FRET signal, demonstrating assay specificity. As additional controls, amyloid precursor protein (APP) co-expression, a known BACE1 substrate (Benilova et al., 2012), reduced FRET efficiency, presumably by competing for the inhibitor probe, while KCNE1 co-expression had no impact on FRET.

Co-expression of KCNQ1 significantly reduced FRET efficiency. A comparable decrease was observed upon co-expression of KCNQ1 with KCNE1 (Fig. 7F). Together with the BiFC assay, these findings support a model in which KCNQ1 selectively impairs BACE1 homomeric multimerization, independent of whether KCNE1 is present or not.

## Discussion

We previously demonstrated that BACE1 expression markedly alters the gating kinetics of KCNQ1 potassium channels, an effect reproduced by the catalytically inactive BACE1 D289N mutant and therefore consistent with a non-enzymatic, physical mechanism (Agsten et al., 2015). Building on these findings, we combined BiFC, single-molecule fluorescence, and electrophysiology to corroborate those results and to map the sites that enabled the physical interactions between KCNQ1, KCNE1, and BACE1.

Figure 8 summarizes the central findings. Each KCNQ1 channel complex predominantly accommodates two BACE1 molecules. Although BACE1 is mainly detected as dimers or trimers at the plasma membrane (Liebsch et al., 2017; Schmechel et al., 2004; Stockinger et al., 2024; Westmeyer et al., 2004), our data suggest that the channel preferentially engages monomeric BACE1. Notably, BACE1 binding and KCNE1 association do not appear to interfere with each other. This conclusion is supported by three observations: (i) BACE1 co-expression does not reverse the KCNE1-induced rightward shift in the voltage dependence of KCNQ1 activation (Fig. 5G); (ii) the effects of BACE1 and KCNE1 are additive; and (iii) KCNQ1 co-expression disrupts BACE1 homomeric assembly irrespective of KCNE1 expression. Together, these findings are consistent with BACE1 and KCNE1 occupying distinct, non-overlapping sites within the KCNQ1 complex and support a model in which BACE1 directly associates with KCNQ1 independently of its protease activity, without affecting KCNE1 binding.

**Figure 8.**
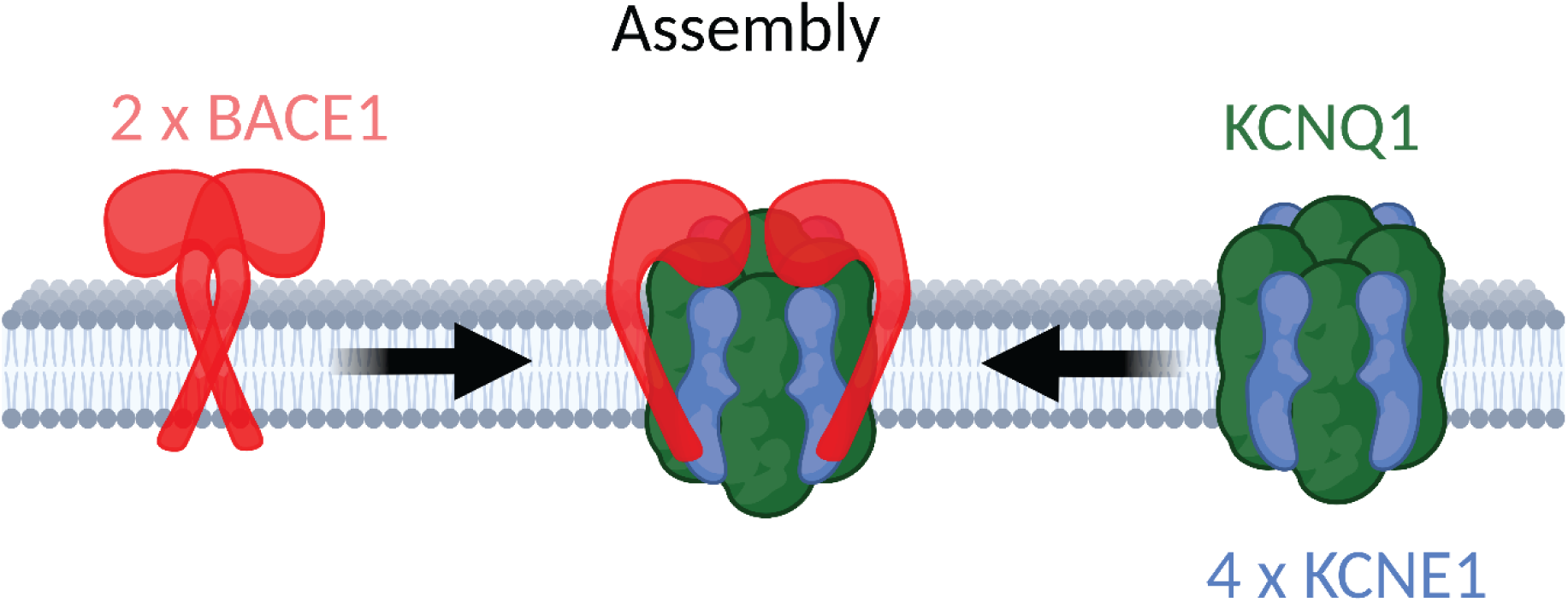
Diagram illustrating the assembly of the KCNQ1–KCNE1–BACE1 channel complex. The channel complex is composed of a homotetrameric KCNQ1 α-subunit together with up to four KCNE1 β-subunits (Murray et al., 2016). Our data indicate that up to two BACE1 molecules can be incorporated into the complex. Notably, although BACE1 typically exists as a homodimer or homotrimer (Stockinger et al., 2024), it associates with the channel as a monomer. The transmembrane segment of KCNE1 appears to be the primary determinant of *I_Ks_* modulation, with additional contributions from its intracellular C-terminus. In contrast, BACE1 engages the channel predominantly via its extracellular catalytic domain. Our findings further suggest that KCNE1 and BACE1 bind to the channel independently.

In agreement with previous reports (Melman et al., 2001; Tapper & George, 2000), we found that the transmembrane segment of KCNE1 is indispensable for its interaction with KCNQ1 α-subunits and for reconstituting an *I_Ks_*-like current in our electrophysiological recordings. Consistent with Tapper & George (2000), the short extracellular N-terminus of KCNE1 is dispensable: our BEE chimera displayed the same voltage dependence of activation as wild-type KCNE1. Replacing the intracellular C-terminus with that of BACE1 (BEB and EEB constructs) still produced an *I_Ks_*-like current, although activation was shifted substantially toward more positive potentials. This phenotype closely parallels that reported for KCNE1 Δ70 and KCNE1 D76N (Chen et al., 2009), in which the absence of the C-terminus was proposed to stabilize the closed state, thereby reducing the probability of channel opening.

Beyond KCNQ1, we previously showed that BACE1 non-enzymatically interacts also with neuronal KCNQ channels (KCNQ2–KCNQ5) (Hessler et al., 2015). This finding was recently corroborated by Dai (2022), who demonstrated that palmitoylation of four C-terminal cysteines in BACE1 recruits KCNQ2/3 channels to lipid-raft microdomains, likely modulating their gating indirectly. In our KCNQ1 experiments, however, BACE1’s effects persisted even when these cysteines were replaced by KCNE1 in the BEE chimera. Moreover, we observed additive effects of BACE1 and KCNE1 on KCNQ1 gating, and the BEE chimera reproduced both (Fig. 5). Together, these findings make it unlikely that BACE1’s influence on KCNQ1 is mediated solely through microdomain co-trafficking. Instead, they support a model in which BACE1 directly associates with KCNQ1 and modulates its gating via its extracellular domain.

Two of the three chimeric constructs containing the extracellular domain of BACE1 (BBE and BEE) produced gating effects on KCNQ1 resembling those of wild-type BACE1 (Figs. 4, 5). The third construct (BEB) could not be reliably assessed because of the pronounced positive shift in activation (Fig. 5E). Interestingly, although the extracellular domain of BACE1 combined with the transmembrane and intracellular regions of KCNE1 can engage the channel and alter its gating, the physical association measured by the BiFC assay was weak and approached background levels (Fig. 3). One plausible explanation is that BACE1 and KCNE1 bind to KCNQ1 at distinct sites and/or with different orientations, preventing optimal alignment of the BiFC constructs. This interpretation is consistent with the additive gating effects of the wild-type proteins, which point to non-overlapping binding sites. We further speculate that the bulky extracellular domain of BACE1 may limit the number of molecules that can bind to a KCNQ1 tetramer, thereby explaining why only two monomeric BACE1 molecules appear to associate with the channel.

Several questions remain open. First, up to four KCNE1 subunits (Murray et al., 2016) are thought to occupy the grooves between subunits in the tetrameric KCNQ1 complex (Sun & MacKinnon, 2020). If all these grooves are occupied, the spatial location of the BACE1 transmembrane domain remains uncertain. Second, it is yet to be determined whether the binding model proposed here also applies to neuronal KCNQ channels, or whether, in that setting, BACE1 modulates gating indirectly by co-trafficking with KCNQ channels to specific membrane microdomains in a palmitoylation-dependent manner (Dai, 2022).

In summary, we propose a revised interaction model for KCNQ1, KCNE1, and BACE1 that posits a direct, non-proteolytic association among all three proteins. The gating effects on KCNQ1 are mediated independently, by BACE1 via its extracellular interface and by KCNE1 via its transmembrane and intracellular interfaces (Fig. 8), allowing for the possibility of separate regulatory pathways. In our previous study, we showed that in the absence of ATP, *I_Ks_* is considerably enhanced by BACE1 (Agsten et al., 2015). Thus, an intriguing aspect of BACE1’s role is its potential to couple *I_Ks_* to the cell’s metabolic state through ATP-dependent effects.

## Acknowledgments

We are grateful to Iwona Izydorczyk for technical assistance. We thank Andreas Gießl for providing BiFC constructs and are grateful for his support. This study was supported by the Studienstiftung des deutschen Volkes to S.K and the Deutsche Forschungsgemeinschaft (DFG, German Research Foundation, HU 2358/1-1) to TH.

